# Parallel Encoding of Speech in Human Frontal and Temporal Lobes

**DOI:** 10.1101/2024.03.19.585648

**Authors:** Patrick W. Hullett, Matthew K. Leonard, Maria Luisa Gorno-Tempini, Maria Luisa Mandelli, Edward F. Chang

## Abstract

Models of speech perception are centered around a hierarchy in which auditory representations in the thalamus propagate to primary auditory cortex, then to the lateral temporal cortex, and finally through dorsal and ventral pathways to sites in the frontal lobe. However, evidence for short latency speech responses and low-level spectrotemporal representations in frontal cortex raises the question of whether speech-evoked activity in frontal cortex strictly reflects downstream processing from lateral temporal cortex or whether there are direct parallel pathways from the thalamus or primary auditory cortex to the frontal lobe that supplement the traditional hierarchical architecture. Here, we used high-density direct cortical recordings, high-resolution diffusion tractography, and hemodynamic functional connectivity to evaluate for evidence of direct parallel inputs to frontal cortex from low-level areas. We found that neural populations in the frontal lobe show speech-evoked responses that are synchronous or occur earlier than responses in the lateral temporal cortex. These short latency frontal lobe neural populations encode spectrotemporal speech content indistinguishable from spectrotemporal encoding patterns observed in the lateral temporal lobe, suggesting parallel auditory speech representations reaching temporal and frontal cortex simultaneously. This is further supported by white matter tractography and functional connectivity patterns that connect the auditory nucleus of the thalamus (medial geniculate body) and the primary auditory cortex to the frontal lobe. Together, these results support the existence of a robust pathway of parallel inputs from low-level auditory areas to frontal lobe targets and illustrate long-range parallel architecture that works alongside the classical hierarchical speech network model.

## INTRODUCTION

To understand how the brain performs remarkable computational feats like speech perception, it is necessary to understand the cortical regions involved and the overall network architecture that connects them. The speech network in the human brain is postulated to have a largely hierarchical organization with information flow from subcortical structures to primary auditory cortex, then to higher order areas on lateral temporal cortex, and finally through dorsal and ventral pathways to apical targets in the frontal lobe^1,2^. While the precise role of these frontal lobe areas in speech perception is debated ^3–7^, a common feature in speech models is frontal lobe inputs arising from lateral temporal cortex^1,2^. Thus, activity in frontal areas during speech perception reflects downstream computational processes inherited from areas like the lateral superior temporal gyrus (STG, **Fig. 1A**, top panel).

**Figure 1.**
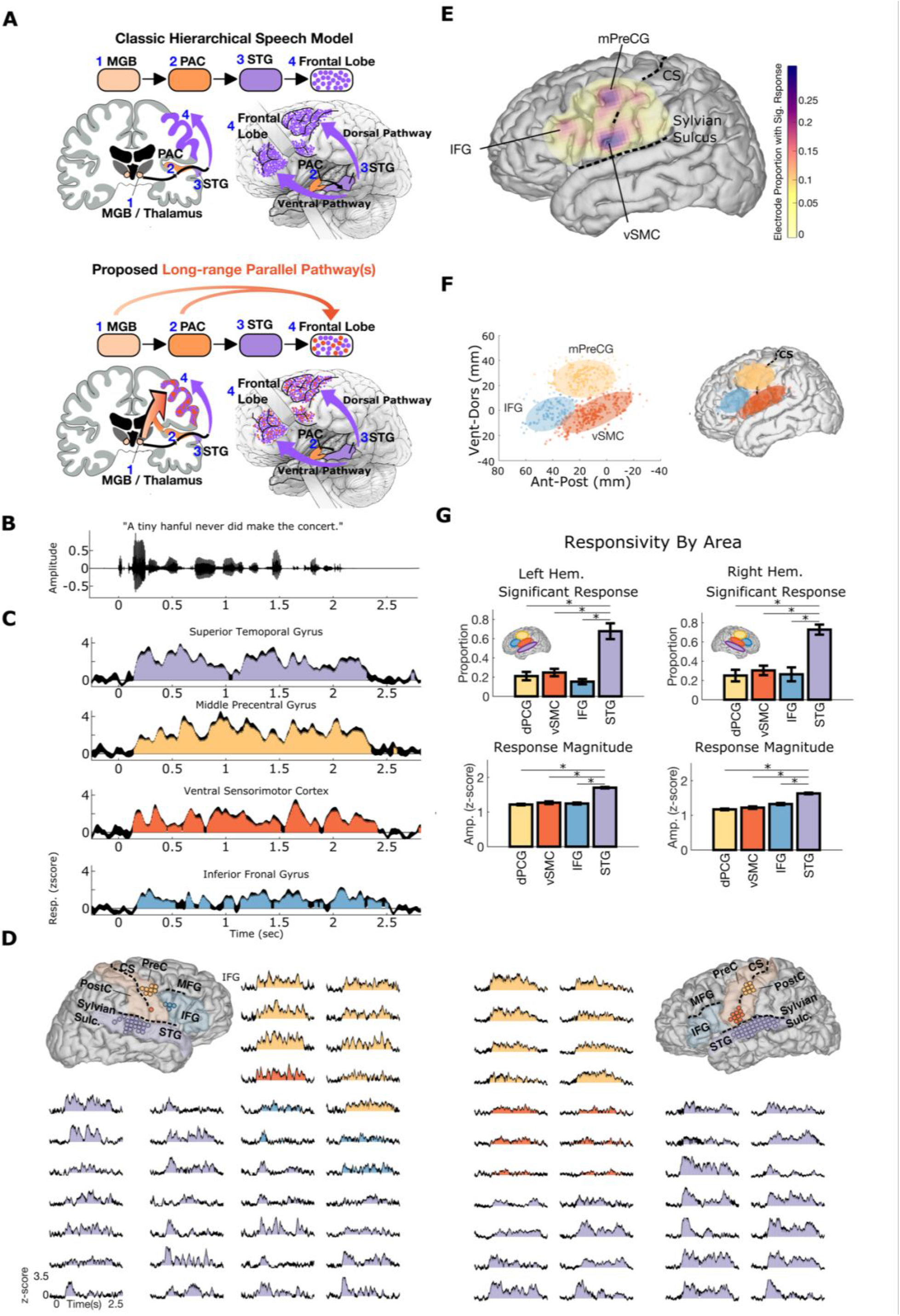
Suprasylvian Speech Response Characteristics. **(A)** Top panel: classic hierarchical speech pathway to prefrontal cortex (1 to 2 to 3 to 4). Bottom panel: additional hypothesized long-range parallel pathways (bright orange arrows from areas 1 and 2 to 4) with inputs from the medial geniculate body (MGB) and primary auditory cortex (PAC) to the frontal lobe. **(B)** Acoustic waveform of a single example sentence presented to participants. **(C)** Example responses to a single sentence from the right hemisphere participant in D. Responses in frontal cortex are robust, have similar onset times, and have comparable temporal variation in magnitude over the course of the sentence. The coloring reflects responses from putative spatial clusters in frontal cortex that are confirmed in E and F. **(D)** Representative example of two participants showing example responses in suprasylvian cortex and temporal cortex to the single sentence shown in B. The brain reconstructions show the spatial locations of responses to this sentence. This shows that the response features described in B are generally present throughout suprasylvian cortex and STG. **(E)** Spatial distribution of significant responses to speech across all participants (significant responses defined as electrodes with p < 0.05 for any response time bin, Bonferroni corrected for the number of time bins in each sentence, Wilcoxon rank-sum test). This shows three areas of peak response probability in IFG, middle precentral gyrus (mPreCG), and ventral sensorimotor cortex (vSMC) (electrodes collapsed to the left hemisphere) **(F)**. Spatial clustering using a Gaussian mixture model demonstrates three areas of speech responses in IFG, mPreCG, and vSMC. The number of clusters was determined by Bayesian information criterion and silhouette criterion values (both converged on 3 clusters)^18^. **(G)** Suprasylvian area versus STG responsivity. The proportion of electrodes with significant responses and the mean response amplitude is lower in IFG, mPreCG, and vSMC than in STG (p < 0.005, two-sided Wilcoxon rank-sum test, Bonferroni corrected). Each area has no left-right hemispheric differences regarding the proportion or magnitude of significant responses (p > 0.05, two-sided Wilcoxon rank-sum test). CS = central sulcus, Hem = hemisphere, IFG = inferior frontal gyrus, MGB = medial geniculate body, mPreCG = middle precentral gyrus, PostC = postcentral gyrus, PreC = precentral gyrus, STG = superior temporal gyrus, Sulc. = sulcus, vSMC = ventral sensorimotor cortex.

However, as our understanding of network architecture in the human brain expands, there is increasing evidence for highly parallelized structure, including in auditory networks^8,9^. Specifically, rather than strictly lateral temporal cortex inputs to the frontal lobe, there may be direct parallel inputs to frontal cortex from low-level areas like primary auditory cortex or the medial geniculate body in the thalamus (**Fig. 1A**, bottom panel, orange arrows). In support of this, studies in primates have found evidence for frontal lobe auditory responses that have short latencies in the range of those observed in lateral temporal cortex^10^ and auditory activity in these areas is well-explained by spectrotemporal features similar to tuning in low-level areas ^11,12^. Additionally, in macaques, areas lateral to primary auditory cortex have direct projections to the frontal lobe, and in humans, electrical stimulation of primary auditory cortex elicits short latency responses in the inferior frontal gyrus^13^. Together, these data suggest a subset of frontal cortex responses may reflect parallel inputs from subcortical nuclei and primary auditory cortex, forming a parallel pathway to frontal cortex that works alongside the classical dorsal-ventral pathway from lateral temporal cortex.

To test this hypothesis, we used electrocorticography (ECoG) to identify speech responses in the frontal and temporal lobes in awake participants while they passively listened to natural speech. We specifically asked whether there is evidence for: (1) short latency speech-evoked responses that are the same in the frontal lobe and temporal lobes, and (2) whether such activity is explained by spectrotemporal encoding models as would be expected for direct inputs from low-level areas such as the primary auditory cortex or the medial geniculate body of the thalamus. Here, we identify three areas in frontal lobe cortex that are activated by speech with latencies that occur simultaneously or even before the earliest latencies in STG. Additionally, neural representations in these short latency populations reflect spectrotemporal content that is essentially indistinguishable from populations in STG. Finally, to further establish evidence for anatomic connections that reflect parallel inputs from lower-order areas to the frontal lobe, we use white matter tractography and functional connectivity and find connections between frontal areas and the medial geniculate body of the thalamus as well as the primary auditory cortex. Overall, these results demonstrate a fundamental divergence in the hierarchical architecture that is usually assumed to underlie cortical speech processing and shows multiple lines of evidence for long-range parallel inputs from low-level areas to frontal lobe cortical regions.

## RESULTS

### Frontal lobe areas are activated by passive speech listening

To characterize the extent to which neural populations in frontal lobe respond to speech, ECoG participants passively listened to 10–40 minutes of natural speech, which consisted of prerecorded sentences (2–4 sec. each) from the phonetically transcribed TIMIT speech corpus^14^.We extracted activity in the high-gamma (70–150 Hz) range^15^, which correlates with spiking activity ^16^ and spike-based tuning properties in the midlaminar auditory cortex^17^. At single electrodes, average responses to an example sentence (**Fig. 1B**) were highly robust and extended through the duration of the sentence in STG (**Fig. 1C**; purple) and three areas in the frontal lobe including middle precentral gyrus (mPreCG; **Fig. 1C**; yellow), ventral sensorimotor cortex (vSMC; **Fig. 1C**; orange), and inferior frontal gyrus (IFG; **Fig. 1C**; blue). Figure 1D shows two example participants that illustrate typical electrode coverage in the left and right hemispheres and responses to a single sentence of speech that are qualitatively similar to responses in STG.

Across all 17 participants, we quantified the spatial distribution of responses to speech in frontal and parietal cortex. We observed significant responses throughout the areas covered by ECoG grids, with three peaks of responsiveness in mPreCG, vSMC, and IFG (**Fig. 1E)**. To further test whether these three peaks reflect spatially clustered responses, we used Gaussian mixture modeling to cluster the electrodes. The three areas that emerged reflect clusters of electrodes in mPreCG, vSMC, and IFG (**Fig. 1F**; cluster number = 3 determined using both Bayesian information criterion, p < 0.05, and Silhouette criterion p < 0.05, Kaufman and Rousseeuw, 1990). Thus, in addition to STG, there are three regions within frontal lobe cortex that have significant responses to natural speech.

We compared the basic response properties in these three suprasylvian areas to STG, which is known to be a critical area that is central to speech perception ^19,20^. To characterize the overall responsivity, we first quantified the proportion of all electrodes within an area that showed a significant response to speech (**Fig. 1G**, top row). Although all three regions had electrodes with significant speech-evoked activity in both hemispheres, there was a lower proportion of speech-responsive electrodes in frontal lobe areas compared to the proportion of speech-responsive electrodes in STG (p < 0.005, two-sided <u>Wilcoxon rank-sum test,</u> Bonferroni corrected; **Fig. 1G**, top row). Additionally, while significantly greater than zero, the average response magnitude in suprasylvian areas was lower than STG (p < 0.005, two-sided <u>Wilcoxon rank-sum test,</u> Bonferroni corrected; **Fig. 1G**, bottom row). For both the proportion of significant electrodes and the response magnitude, there were no significant differences between hemispheres (p > 0.05; two-sided [Wilcoxon rank-sum test](https://www.mathworks.com/help/stats/ranksum.html#bti4z5t)), suggesting that frontal lobe speech-evoked activity is similar between right and left hemispheres^21,22^.

### Frontal lobe areas have short-onset latencies like the superior temporal gyrus

Having established the existence of robust speech-evoked activity throughout bilateral frontal lobe cortex during passive listening, we asked whether any of these neural populations had short-onset latencies that are synchronous or faster than the earliest onset latencies in lateral temporal cortex. If present, this would be consistent with parallel pathways from lower-order auditory areas to frontal and temporal lobes. Direct visual examination of speech evoked activity showed near synchronous shortest latency responses across frontal areas and STG (**Fig. 2B**). Furthermore, there were within-participant examples of frontal electrodes with onset latencies synchronous or earlier than the shortest onset latencies in the temporal lobe (**Fig. 2C**). To quantify this, we calculated the response onset latencies across all participants similar to prior characterizations in humans and primates **(Fig. 2A, 2D**, Camalier et al., 2012; Nourski et al., 2014). This showed electrodes throughout temporal, frontal, and parietal lobes with short response latencies, often less than 80ms. For each region, we quantified the latency distribution, which showed similar short latency responses across all areas (**Fig. 2D**). To test for significant differences in short-onset latencies between areas, we used a temporal cutoff to define the left-sided tail of each distribution as the short-onset latencies of interest (**Fig. 2E**). The temporal cutoff ranged from 80-720ms, with significance testing at each cutoff. As shown, onset latencies were not significantly different between frontal lobe areas (IFG, mPreCG, vSMC, and STG) and STG for latencies up to 200ms in the left hemisphere and 240ms in the right hemisphere (**Fig. 2E**, p > 0.05, two-sample Kolmogorov-Smirnov test, Bonferroni corrected). This data shows short-onset latencies within the same hemisphere are not significantly different between frontal lobe areas and STG. Thus, in response to natural speech, IFG, mPreCG, and vSMC are active beginning at the same time as neural populations in STG, consistent with parallel inputs to STG and frontal lobe areas.

**Figure 2.**
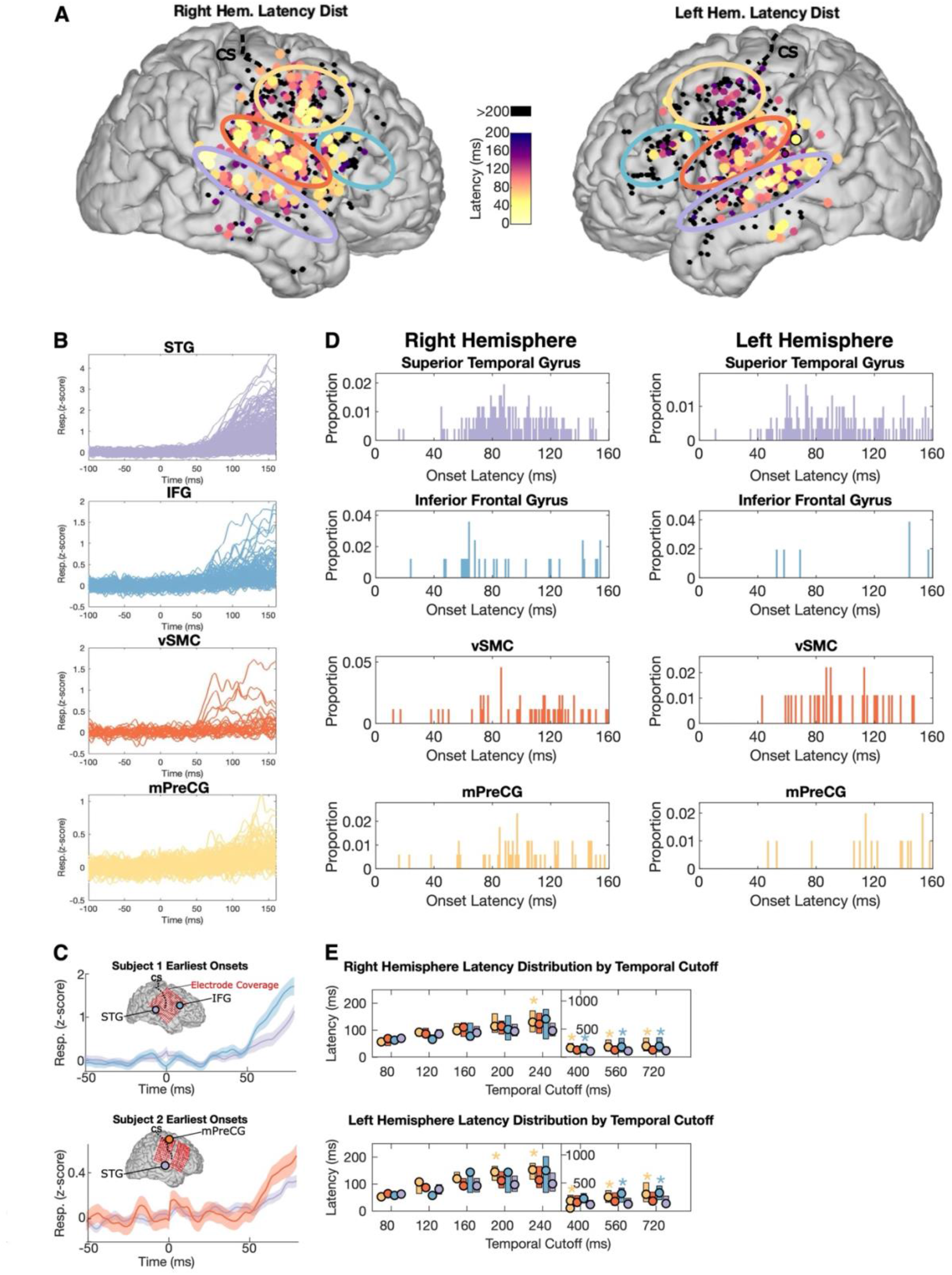
Latency Characterization Across all Participants. **(A)** Spatial map of response latencies show short-onset latencies in STG, IFG, mPreCG and vSMC (onset defined as first 1-ms time bin in which the response p value is < 0.05 for 15 consecutive 1-ms bins; Wilcoxon rank-sum test; Camalier et al., 2012; Nourski et al., 2014) **(B)** All responses across participants with onset latencies less than 200ms. **(C)** Two example participants showing the electrode sites with shortest onset latency in the STG and frontal lobe. As shown, the earliest onset latencies within a subject are similar between the frontal lobe and STG. **(D)** Latency distribution by area for onset latencies less than 160ms (the proportion y-axis is relative to the full distribution of latencies up to 1600ms). This shows the earliest onset latencies are similar across areas and is consistent with low-latency parallel inputs to each area. **(E)** Latency distribution boxplots (median + interquartile range) for all onset latencies up to the temporal cutoff specified on the x-axis (80 to 720ms). There is no significant difference between STG and IFG, mPreCG, or vSMC up to 240ms in the right hemisphere and 200ms in the left hemisphere (* = p < 0.05, two-sample Kolmogorov-Smirnov test, Bonferroni corrected). These indistinguishable short-onset latency times are consistent with a subset of parallel inputs to frontal cortex and STG rather than strictly hierarchical inputs from STG to frontal cortex. CS = central sulcus, IFG = inferior frontal gyrus, mPreCG = middle precentral gyrus, STG = superior temporal gyrus, vSMC = ventral sensorimotor cortex.

### Short-onset latency inputs to the frontal and temporal lobe encode the same spectrotemporal speech information

A feature of most functional-anatomical models of speech is frontal lobe areas receive inputs from the temporal lobe through dorsal and ventral pathways and may reflect qualitatively different (and perhaps higher-order) representations of speech ^1,2^.Given the existence of parallel short latency responses in the frontal and temporal lobes, we asked if the spectrotemporal information encoded in these responses fundamentally differed between lobes. To test this, we computed spectrotemporal receptive fields (STRFs, **Figure 3A**) for each site within STG, IFG, mPreCG, and vSMC for short-onset latency sites (< 200ms). In all three frontal lobe areas, responses to speech were well-predicted by STRFs (mean r = 0.39 ± .24, **Fig. 3B**), with some electrodes reaching r=0.75, demonstrating that frontal lobe neural population activity is well-explained by spectrotemporal models.

**Figure 3.**
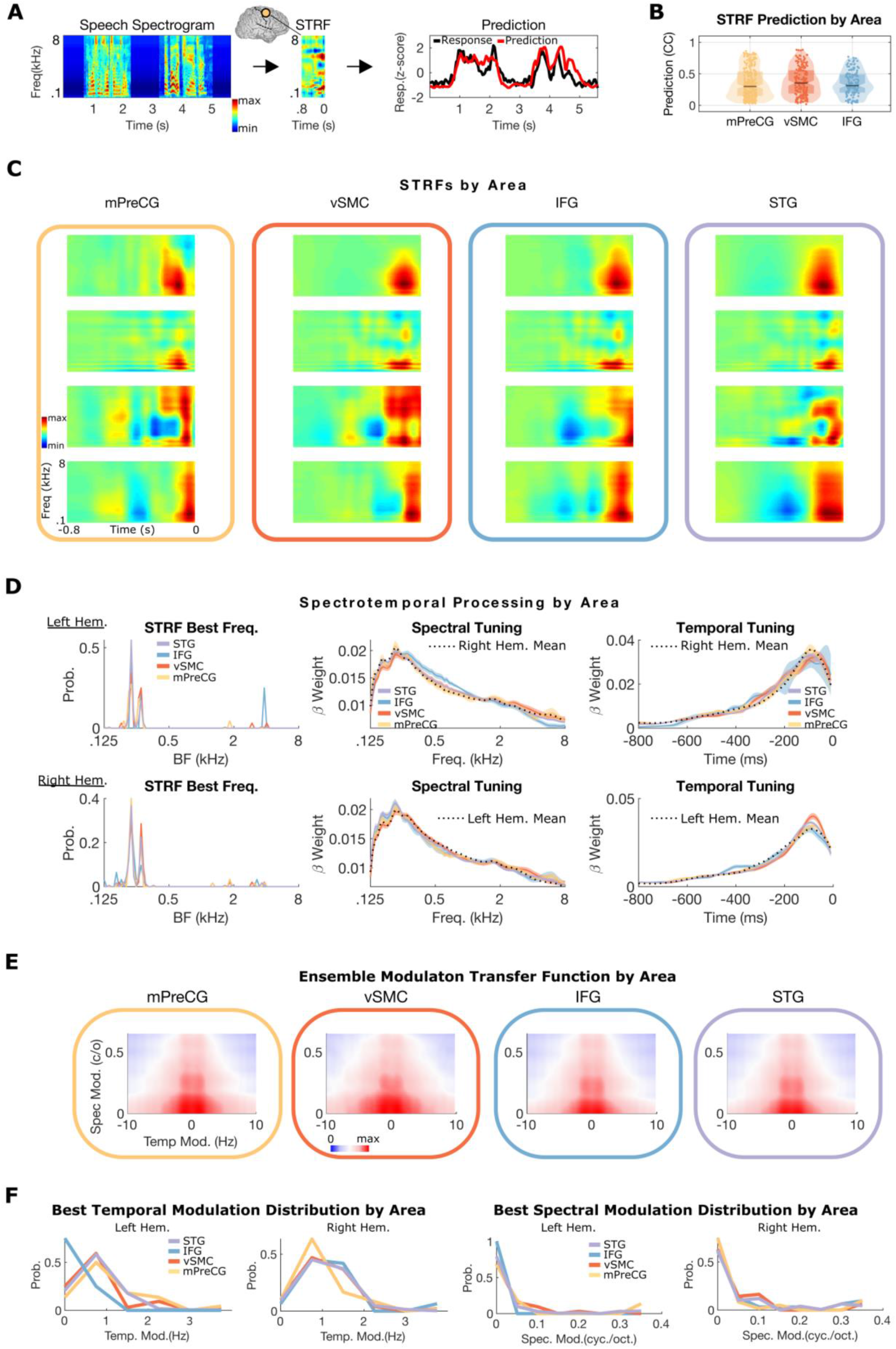
Sites with Short-onset Latencies in Frontal Cortex Encode the Same Spectrotemporal Information as STG. **(A)** Example middle precentral gyrus STRF with the predicted and actual response to a single sentence. Predicted responses are obtained by convolving the spectrogram with the STRF and are proportional to the similarity between the stimulus’s spectrotemporal content and the STRF structure. **(B)** STRF Prediction value distributions for all electrodes with a significant response to speech in mPreCG, VSMC, and IFG (suprasylvian cortex global mean Pearson correlation coefficient 0.39 ± .24; tested on held-out data). STRFs, which are predictive of neural responses, characterize spectrotemporal encoding at each site and are consistent with the presence of spectrotemporal representations in these areas. **(C)** Examples showing the high degree of STRF similarity between suprasylvian areas and STG. **(D)** There is no significant difference in short-onset (< 200 ms) STRF tuning parameters across suprasylvian sites and STG, including: best frequency tuning (p > 0.05 Kruskal-Wallis test), spectral, or temporal tuning (p > 0.05 cosine similarity permutation test of mean spectral and temporal tuning vectors). Spectral and temporal tuning plots show the mean ± s.e.m for each area in each hemisphere; the total contralateral hemisphere mean tuning is plotted as a dotted line for reference. **(E)** Ensemble modulation transfer functions by area. The average modulation tuning for each area has a high degree of similarity. **(F)** There is no significant difference in temporal or spectral modulation tuning between frontal lobe areas and STG (p > 0.05 cosine similarity permutation test of mean spectral and temporal modulation tuning vectors^30^). BF = best frequency, CC = Pearson correlation coefficient, c/o = cycles per octave, Hem. = hemisphere, IFG = inferior frontal gyrus, mPreCG = middle precentral gyrus, Spec, Mod. = spectral modulation (cycles, octave), STRF = spectrotemporal receptive field, vSMC = ventral sensorimotor cortex.

Next, we asked whether the spectrotemporal content encoded in short-onset (< 200ms) frontal lobe electrodes is similar to STG. Based on visual examination of short-onset STRFs from each frontal lobe area and STG, there was a high degree of similarity in spectrotemporal encoding (**Fig. 3C**). To quantify this we examined the STRF weights and computed three key parameters (**Fig. 3D**): (1) the best frequency (first column) ^25,26^, (2) spectral tuning (second column) ^27,28^, and (3) temporal tuning (third column) ^27,29^. For each of these parameters, there was no significant difference across regions (p > 0.05 Kruskal-Wallis test and cosine similarity permutation test of mean spectral and temporal tuning vectors^30^).

While individual STRF tuning properties were not different between STG and frontal lobe areas, the *joint* combination of these tuning properties in each STRF characterizes the spectrotemporal processing at each site. To jointly characterize spectrotemporal processing for each electrode across regions, we transformed the STRFs into their modulation transfer functions (MTFs) ^31–33^. MTFs are the modulation domain representation of spectrotemporal processing characterized by STRFs ^32,34^. They summarize neural processing in terms of spectral and temporal modulation tuning and are frequently used to characterize processing within the auditory system ^35–40^. As shown in figure 3E, the ensemble (mean) modulation transfer function across frontal lobe areas and STG is similar. To quantify this, the mean temporal modulation tuning, and spectral modulation tuning distributions derived from the MTFs were calculated (**Fig. 3F**) and no significant differences in temporal or spectral modulation tuning were observed across frontal lobe areas and STG (p > 0.05 cosine similarity permutation test of mean spectral and temporal modulation tuning vectors^30^). Together, these results indicate that the short latency responses in frontal lobe areas and STG encode the same spectrotemporal representations of speech, consistent with parallel spectrotemporal speech representations in the temporal and frontal lobes.

While short-onset (< 200 ms) electrodes in the frontal lobe are well-predicted by spectrotemporal speech representations, it is possible that they reflect higher-order linguistic features that are correlated with the speech spectrogram. To test this alternative, we compared encoding performance for STRFs to semantic-based encoding models generated with word vector representations of speech derived from the FASTTEXT data set ^41^. We expected that both spectrotemporal and semantic encoding models would perform well in each region, however, they would differ from each other as a function of short or long onset latency. Specifically, we hypothesized that the STRF would be a better model for electrodes with short-onset latencies since these populations reflect direct inputs from low-level areas such as the medial geniculate body or primary auditory cortex.

Confirming our hypothesis, we found that electrodes with short-onset latencies (< 200 ms) in IFG, mPreCG, and vSMC show significantly higher STRF model predictions than semantic model predictions, consistent with dominant spectrotemporal representations at those sites **(Fig. 4A**, p < 0.01, Wilcoxon rank-sum test). At longer latencies (> 200ms), spectrotemporal and semantic encoding models were not significantly different, further supporting the notion that these short latency electrodes reflect spectrotemporal representations. Overall, these data demonstrate that neural populations in the frontal lobe with short-onset responses encode spectrotemporal representations of sound similar to populations in more low-level auditory areas and STG.

**Figure 4.**
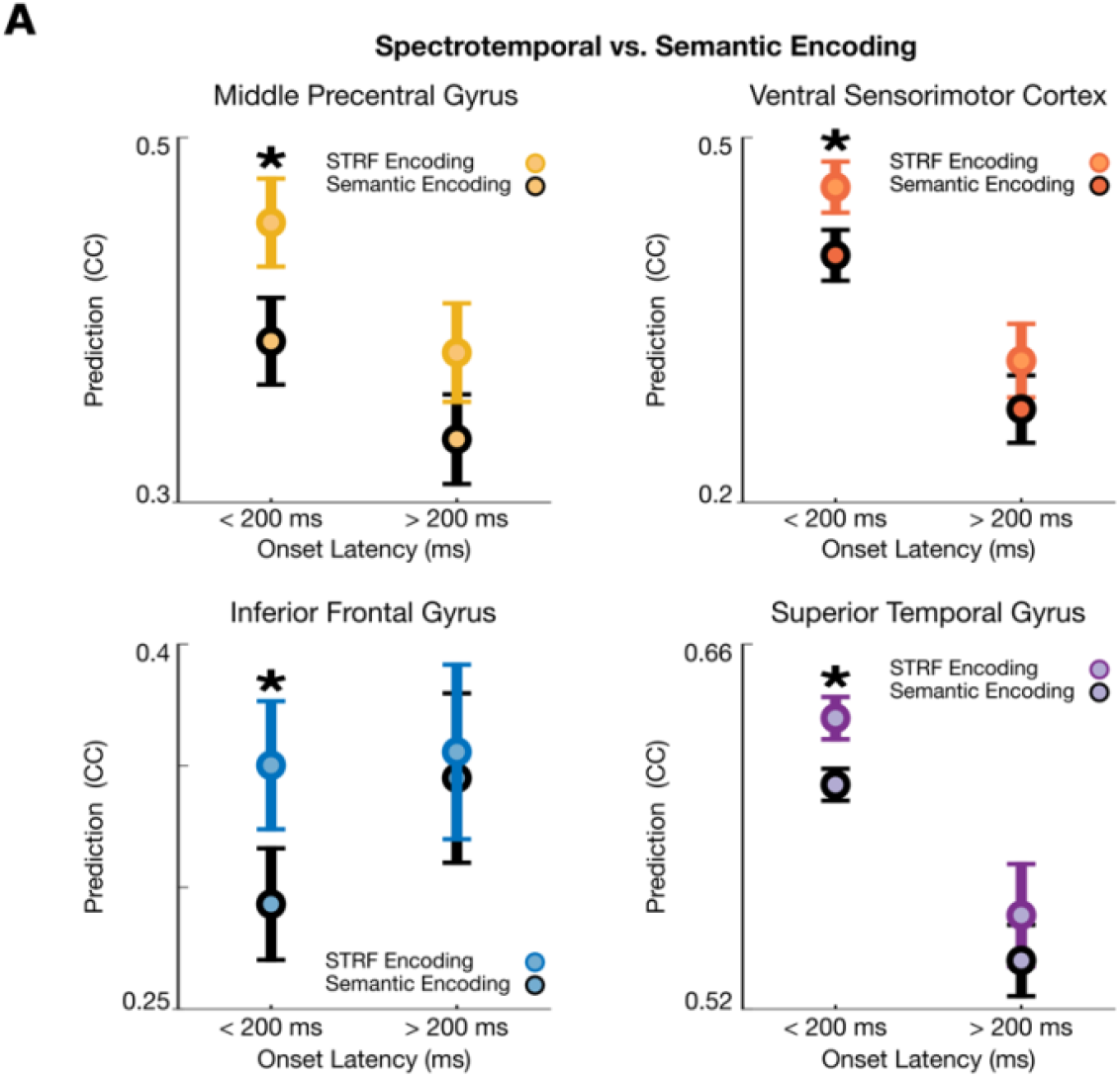
Spectrotemporal Encoding Models are Better than Semantic Encoding Models for Short-onset Latency Sites in Frontal and Temporal Lobes. **(A)** Short-onset latency sites in each area encode low-level spectrotemporal information. Sites with onset latencies less than 200ms show significantly higher STRF model predictions than semantic model predictions consistent with dominant spectrotemporal over semantic coding at these short-onset latency sites. This is consistent with short-onset latescent sites encoding spectrotemporal speech information in parallel (p < 0.001, Wilcoxon rank-sum test). Longer onset latency sites in the same areas show no difference between spectrotemporal or semantic encoding models. CC = Pearson correlation coefficient, STRF = spectrotemporal receptive field.

### White matter tractography and functional connectivity show connections from thalamus and primary auditory cortex to frontal lobe areas

To establish whether the short latency responses with STG-like spectrotemporal encoding observed in frontal lobe cortex reflects a direct anatomical pathway from lower-level auditory areas, we used diffusion tensor imaging (DTI) tractography. Based on the known connectivity patterns with STG, we hypothesized that the medial geniculate body (MGB) within the thalamus and the primary auditory cortex within Heschl’s gyrus are key areas that might have parallel projections to frontal lobe cortex.

Using data from 842 individuals from the Human Connectome project ^42^, we calculated white matter tractography in each hemisphere with the MGB and Heschl’s gyrus as seed regions. To localize the MGB, we used MNI coordinates in conjunction with the terminal point of white matter tracts from the inferior colliculus to the MGB within the thalamus (Fig. 5A). We restricted the suprasylvian target region of interest to the total area spanned by IFG and ventral to mid-Rolandic cortex. We identified tracts from MGB that projected to IFG, mPreCG, and right vMSC (**Fig. 5A**). We also identified tracts from Heschl’s gyrus that projected to vSMC and left IFG.

**Figure 5.**
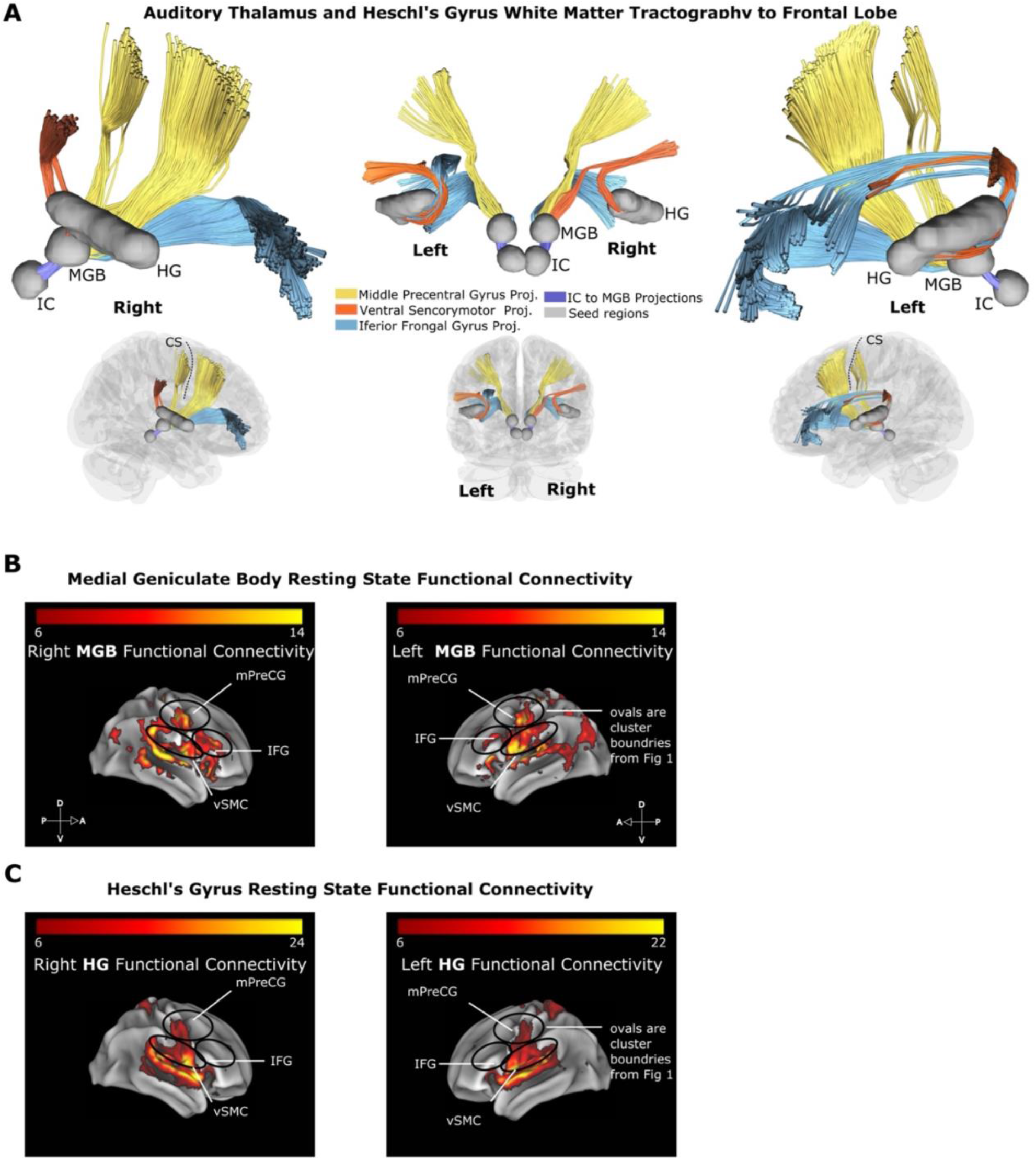
Auditory Thalamus and Heschl’s Gyrus White Matter Tractography and Functional Connectivity with Frontal Lobe. **(A)** White matter tractography between the medial geniculate body (MGB), Heschl’s gyrus (HG), and suprasylvian cortex (IFG, middle precentral gyrus (mPreCG), and ventral sensorimotor cortex (vSMC)). White matter tracts are color-coded by the suprasylvian site of termination. This analysis demonstrates tractography data consistent with direct MGB-to-frontal cortex and HG-to-frontal cortex white matter tracts. **(B)** Resting-state functional connectivity of the medial geniculate body and **(C)** Heschl’s gyrus showing functional connectivity to frontal lobe areas middle precentral gyrus, ventral sensorimotor cortex, and inferior frontal gyrus (p < 5 × 10^−5^ after peak-level family wise error (FWE) correction for multiple comparisons). Overall, these data provide additional structural and functional evidence for parallel pathways to frontal lobe areas from low-level areas such as Heschl’s gyrus and the medial geniculate body. FC = functional connectivity; HG = Heschl’s gyrus; IFG= inferior frontal gyrus; IC = inferior colliculus, MGB = medial geniculate body, mPreCG = middle precentral gyrus, vSMC = ventral sensorimotor cortex.

To test whether identified tracts relate to functional measures, we also calculated resting-state functional connectivity between the medial geniculate body or Heschl’s gyrus to all of cortex using fMRI. For the MGB seed region, there was significant functional connectivity with IFG, mPreCG, and vSMC (**Fig. 5B**, p < 5 × 10^−5^ after peak-level family-wise error (FWE) correction for multiple comparisons). For the Heschl’s gyrus seed region, there was significant functional connectivity to mPreCG and vSMC (**Fig. 5C**, p < 5 × 10^−5^ after peak-level family-wise error (FWE) correction for multiple comparisons). Together, white matter tractography and resting-state functional connectivity demonstrate that short latency evoked activity that encodes speech-relevant spectrotemporal content in frontal lobe cortex reflects a parallel auditory pathway that functions alongside the classical hierarchical speech network mediated by STG.

## DISCUSSION

We examined the nature of short latency, speech-evoked activity in frontal lobe areas that are not typically associated with auditory processing. Using high-density ECoG, we found that a subset of neural populations in frontal lobe cortex exhibited response latencies that were synchronous with or preceded the earliest response latencies in STG. Spectrotemporal representations in these populations were largely indistinguishable from those found in STG, encoding key spectral and temporal modulation rates for speech. Finally, we identified white matter tracts that connect early auditory regions like MGB and Heschl’s gyrus with frontal lobe areas and were associated with resting-state functional connectivity. Together, these results indicate the existence of a direct pathway between low-order areas (medial geniculate body, primary auditory cortex) and neural populations in frontal cortex, which work in parallel to the more well-established hierarchical pathway via the temporal lobe.

Sensory systems, including vision and hearing, are known to have hierarchical structure, with dominant feedforward connections from the thalamus to primary sensory cortex to higher-order areas within frontal cortex ^1,43,44^. In addition to this hierarchical structure, locally, within nearby levels, there are short-range parallel connections^9^. However, the presence of long-range parallel connections in humans from low-level areas such as MGB directly to apical areas in frontal cortex have not been well described. The present results demonstrate that, in addition to short-range parallel connections, long-range parallel connections exist from thalamus and primary auditory cortex to areas in the frontal lobe.

It may be surprising to find such long-range connections, particularly as part of a network involved in processing such a complex stimulus as speech. However, three important frontal lobe characteristics suggest a direct, parallel auditory pathway. First, these frontal areas are highly heterogeneous, and sub-populations of neurons in primates appear to be responsive to the acoustic properties of sound, similar to the tuning properties seen in lower-order areas^11^. Second, prior work in primates has suggested the presence of short onset latencies to sound ^10^, and in humans, electrical stimulation of primary auditory cortex elicits short latency responses in the inferior frontal gyrus^13^. Lastly, in rhesus macaques, non-primary auditory areas, directly adjacent to primary auditory cortex, have tracer-defined anatomic projections to the frontal lobe. The present work shows short-latency neural populations with acoustic representations that are indistinguishable from STG consistent with direct parallel transmission of low-level representations of speech directly to frontal cortex and is further supported by structural evidence of white matter tracts connecting these areas.

Regarding the potential function of a parallel pathway, an intriguing possibility is that these frontal lobe auditory populations facilitate the integration of processes that are canonically associated with frontal areas and lower-level auditory^45–48^. For example, there is substantial data supporting the role of areas like IFG and ventral precentral gyrus in top-down modulation during speech perception ^45–47,49,50^. An important aspect of top-down modulation is the remarkable speed with which it occurs; for example, there is behavioral and neural evidence for nearly instantaneous perceptual warping in phenomena like phoneme restoration ^46,51,52^. Whereas most hierarchical models of sensory processing assume that frontal cortex receives inputs from areas like posterior STG, rapid, frontal cortex-mediated perceptual restoration may be facilitated by direct, parallel thalamocortical input to these areas.

Similarly, a parallel auditory pathway to neural populations throughout sensorimotor speech cortex may play a role in real-time acoustic feedback for articulatory control ^53,54^. Neighboring – or possibly even overlapping – neural populations that directly control articulation ^55–60^ show sensory responses during passive listening ^61–63^, which could be used in both speech-motor planning and for real-time feedback from the acoustics generated by articulatory output. In either potential role (top-down modulation or real-time acoustic feedback) this long-range parallel pathway gives frontal lobe cortical areas real-time access to the primary spectrotemporal input signal to the cortex in addition to higher-level representations it receives by other areas such as lateral temporal cortex.

Finally, we demonstrated that short latency, parallel auditory responses in frontal lobe cortex are supported by white matter tracts from the thalamus and Heschl’s gyrus. To our knowledge, these white matter tracts have not been identified previously. This is likely because the most robust analysis requires a specific hypothesis about seed regions and targets, and the specific pairing of areas like MGB and frontal cortex may have yet to be considered. However, it is important to note that for DTI tractography to identify a robust tract, it must be composed of a relatively large, anisotropic bundle of fibers ^64^. Thus, these newly identified tracts are not minor examples of highly specific connections but rather are substantial, long-range myelinated fibers.

We do not disregard the importance of canonical processing pathways, or the fundamental hierarchical nature of the brain^1,2,65^. Indeed, for speech, it is well-established that structures like the arcuate fasciculus support a major pathway between auditory temporal and frontal cognitive and motor regions^66–68^. Instead, we propose that parallel inputs provide additional computational resources that support the rapid processing required for sounds like speech. Further work is necessary to understand how these pathways work together and whether they have distinct targets within frontal cortex (perhaps suggested by the distributed and relatively sparse nature of short latency responses in the present results). However, taken together, the existence of a direct, parallel auditory pathway to frontal cortex supports the notion that the degree to which neural systems underlying speech perception are purely or largely hierarchical needs to be examined more closely.

## ACKNOWLEDGMENTS

We would like to thank participants, members of the Chang lab, and the EEG technologists at the University of California, San Francisco for their time and effort in acquiring electrophysiological data. Research reported in this publication was supported by the National Center for Advancing Translational Sciences of the NIH under Award Number (5TL1TR001871-05 to P.W.H.). This work was supported by grants from The National Institute of Deafness and Other Communication Disorders (R01-DC012379, K24 DC015544) and The National Institute of Neurological Disorders and Stroke (U01-NS117765). This research was also supported by Bill and Susan Oberndorf, the Joan and Sandy Weill Foundation, and the William K. Bowes Foundation.

## AUTHOR CONTRIBUTIONS

Conceptualization, P.W.H, M. K. L., E.F.C; Methodology, P.W.H, M. K. L., M. L. G., M. L. M., and E. F. C; Software, P.W.H. M. L. M.; Formal Analysis, P.W.H and M. L. M.; P. W. H., M. L. K., M. L. M.; Data Curation P. W. H., M. L. M., Writing – Original Draft P. W. H., Writing – Review & Editing Investigation, M.E., A.N.V., N.A.V., S.C.P., and S.Y.W.; Writing – Original Draft, S.C.P. and S.Y.W.; Writing – Review & Editing P.W.H, M. K. L., M. L. G., M. L. M., and E. F. C; Visualization, P. W. H., M. L. K., and M. L. M.; Supervision P. W. H., M. K. L., and E. F. C.; Project Administration P. W. H., M. K. L., and E. F. C.; Funding Acquisition P.W.H, P.W.H, M. K. L., M. L. G., M. L. M., and E. F. C.

## METHODS

### Participants and Neural Recordings

ECoG arrays (interelectrode distance = 4mm) were placed subdurally in 17 patient volunteers (9 right hemisphere, 8 left hemisphere) undergoing a neurosurgical procedure for the treatment of medication-refractory epilepsy. All participants were native English speakers; all were fluent in English. All participants had normal hearing and no communication deficits. All experimental protocols were approved by the University of California, San Francisco Institutional Review Board and Committee on Human Research. The location of array placement was determined by clinical criteria alone. Participants were asked to passively listen to 10 – 40 minutes of natural speech while ECoG signals were recorded simultaneously. Signals were amplified and sampled at 3052 Hz. After the rejection of electrodes with excessive noise or artifacts, signals were referenced to a common average and the high gamma band (70 - 150 Hz) was extracted as the analytic amplitude of the Hilbert transform ^15,69^. Signals were subsequently downsampled to 100 Hz. For onset latency analsysis the high gamma band was also extracted using Morlet wavelet decomposition and downsampled to 1000Hz. The resulting signal for each electrode was z-scored based on the mean and standard deviation of activity during the entire block.

### Stimuli

Speech stimuli were delivered binaurally through free field speakers at approximately 70 dB average sound pressure level. The frequency power spectrum of stimuli spanned 0 - 8000 Hz. The stimulus set consisted of prerecorded (2 – 4 second) sentences from the phonetically transcribed TIMIT speech corpus with one-second silent intervals between each sentence presentation ^14^. To quantify response characteristics to individual sentences for responsivity and onset latency analysis we analyzed responses to ten unique sentences (each unique sentence repeated ten times). Each participant was also presented an additional 115-489 unique, non-repeated sentences for receptive field analysis. The total speech corpus included 286 male and 116 female speakers, with 1-3 sentences spoken per speaker, and unique lexical content for each sentence.

### Responsivity and Spatial Analysis

Ten TIMIT sentences were presented randomly ten times in each participant (100 sentence presentations total). Neural data was aligned by the onset of sound for each sentence. The mean evoked potential to each of the ten sentences was then computed. To test for speech evoked responses, for each time bin of the evoked potential after sound onset, a Wilcoxon rank-sum test was performed to test for a significant difference from baseline (p < 0.05, Bonferroni corrected for the number of sample bins in the sentence). To visualize electrode coordinates in MNI space, we performed nonlinear surface registration using a spherical sulcal-based alignment in Freesurfer, aligning to the cvs_avg35_inMNI152 template^70^. This nonlinear alignment ensures that electrodes on a gyrus in the participant’s native space remain on the same gyrus in the atlas space but does not maintain the geometry of the grid ^71^. For spatial analysis, significant responses were projected to the left hemisphere in all participants, given that there were no significant differences in the proportion of responsive sites in each area or amplitude of responses between the left and right hemispheres in suprasylvian cortex. Spatial clustering was performed using a mixture of gaussians model with the number of clusters (cluster number = 3) identified by both Bayesian information criterion and Silhouette criterion.

### Analysis

#### Onset Latency Analysis

Each trial of the repeated TIMIT sentences was aligned by the onset of sound. The mean evoked potential for each sentence was then computed. The 500ms before sentence onset served as the baseline for comparisons. For each 1-ms bin of the evoked potential after sound onset, a Wilcoxon rank-sum test was performed to test for a significant difference from baseline (p < 0.05). Like prior ECoG response latency work in the human auditory cortex or primate auditory cortex, response latency was defined as the time in which the mean evoked potential was significantly different from baseline and remained significant for 15 ms^23,24^. The shortest sentence onset latency for each electrode was defined as the speech onset latency for that site.

#### Encoding Analysis

STRF and semantic encoding models were fit using normalized reverse correlation (Theunissen et al., 2001) with open-source code available at: http://strfpak.berkeley.edu/. Regularization was controlled by fitting a tolerance hyperparameter via cross-validation^74^. STRFs were computed on an estimation set (90% of the total data) and cross-validated on a test set, which was withheld from the estimation process (10% of the data). For STRF fitting, spectrogram representations of speech stimuli were generated using a cochlear model of auditory processing ^75^. For the semantic encoding model, word vector representations of the TIMIT speech corpus were derived from the FASTTEXT data set ^41^ by mapping each 10ms segment of the speech signal to the corresponding word vector representation for that word.

#### Modulation Tuning

To characterize modulation tuning, the modulation transfer function (MTF) for each site was computed by taking the magnitude of the two-dimensional Fourier transform (𝔍_2_ {⟨}) of each STRF:

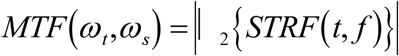

Where (*t, f*) are time, and frequency and (*ω*_*t*_, *ω*_*s*_) are temporal and spectral modulation, respectively^25^.

#### Tractography Analysis

For white matter tractography, we applied a deterministic diffusion fiber tractography algorithm ^76^ using the spin distribution function (SDF) template created by Yeh et al. ^77^ and publicly available diffusion data (http://brain.labsolver.org) with high angular and high spatial resolution from 842 individuals from the Human Connectome Project ^42^. For the tractography analysis, seed regions of interest were specified as the medial geniculate body or Heschl’s gyrus within each hemisphere. A single target region of interest within each hemisphere was specified as the total area spanned by mid and ventral Rolandic cortex, and IFG. Tractography was performed within each intrahemispheric seed-target combination independently. Tractography seed and target regions, with exception of the medial geniculate body, were specified using the Human Connectome Project Multi-Modal Parcellation version 1.0 anatomical atlas and the FreeSurfer Destrieux anatomical atlas ^78–80^. The medial geniculate body seed region coordinates were determined from previously reported MNI coordinates ^81^ and validated by demonstration of positive tractography from the inferior colliculus (the auditory input nucleus to the medial geniculate body) to the full thalamus.

#### Functional Connectivity Analysis

Brain functional images were acquired in a cohort of 50 neurologically intact participants at Siemens 3-Tesla Prisma scanner located at UCSF. We collected 560 T2*-weighted EPI volumes for each individual with the following parameters: TR/TE = 850/32.8 ms, flip angle = 45°, voxel size = 2.2x2.2.x2.2 mm^3^, field-of-view = 211 × 211 mm^2^, multi-band accelerating factor = 6. Image preprocessing consists of slice-time correction, realignment to the mean functional image, assessment for rotational and translational head motion, and correction for susceptibility-induced distortions. Functional images are then normalized to the EPI template in the MNI space with a combination of rigid, affine, and nonlinear warping. After smoothing the images with a 5 mm full width at half maximum (FWHM) Gaussian kernel, CSF and white matter tissue probability maps were then used to compute the mean time series used as regressors. Functional data were then bandpass filtered (0.008 Hz < f < 0.15 Hz), and the nuisance variables were regressed out from the data, which included the six motion parameters, the first derivative and quadratic terms, as well as CSF and white matter time series. Seed ROIs were located bilaterally in the MBG (MNI coordinates: left x=-9, y=-23, z=-1; right x=8, y=-23, z=-1) and in the Heschel’s gyrus. Single-subject correlation maps were generated by calculating the *r*–Pearson correlation coefficient between the average BOLD signal time course from the seed ROIs and the time course from all other voxels of the brain. Finally, correlation maps were converted to *z*-scores, and group-level connectivity maps were calculated for each seed with a statistical threshold at p < 5 × 10^−5^ after peak-level family-wise error (FWE) correction for multiple comparisons.

